# RfaH is Essential for Virulence and Adaptive Responses in *Yersinia pseudotuberculosis* Infection

**DOI:** 10.1101/2025.02.26.640407

**Authors:** Joram Kiriga Waititu, Kristina Nilsson, Gerald Larrouy-Maumus, Tiago R D Costa, Kemal Avican

## Abstract

We previously demonstrated that increased expression of the gene encoding transcriptional antiterminator RfaH during *Yersinia pseudotuberculosis* transcriptional reprogramming necessary for adapting to persistent infection. RfaH is known to regulate expression of the O-antigen biosynthesis operon in *Y. pseudotuberculosis*. In this study, we examined the role of RfaH in virulence, bacterial physiology under infection-relevant stress conditions, and determined the RfaH regulon in *Y. pseudotuberculosis*. We employed a mouse infection model and phenotypic assays to test RfaH’s role in virulence and physiology, as well as RNA sequencing, including O-antigen biosynthesis-deficient strains. Our findings demonstrate that loss of *rfaH* significantly attenuates virulence, reducing the capacity of *Y. pseudotuberculosis* to establish persistent infection. RfaH expression is increased during the stationary growth phase and under various stress conditions, such as high osmolarity and temperature. Functional assays revealed that the *ΔrfaH* strain displayed defects in swimming and increased clumping, indicating altered surface properties affecting motility. Transcriptomic analysis showed that the absence of *rfaH* led to downregulation of genes involved in virulence besides O-antigen biosynthesis operon, suggesting RfaH’s critical role in virulence and host adaptation. Notably, we identified a hypothetical non-coding RNA encoded within the 5’-UTR of the O-antigen biosynthesis operon, which may regulate gene expression of the urease operon in *Y. pseudotuberculosis*. Collectively, our findings suggest that RfaH is essential for the virulence and adaptive capacity of *Y. pseudotuberculosis* to colonize the host. This study provides insights into regulatory mechanisms that facilitate bacterial survival in hostile environments and highlights the importance of RfaH and its regulatory targets in the pathogenesis of *Y. pseudotuberculosis*.

**Author Summary:** For bacterial pathogens to establish infection and persist in the host, they must adapt to harsh environments and fine-tune gene expression accordingly. The transcriptional antiterminator RfaH plays a pivotal role in regulating key genes essential for adaptation and virulence in *Y. pseudotuberculosis*. In this study, we explored the function of RfaH in bacterial physiology, stress responses, and infection dynamics. Using a mouse infection model, we found that loss of RfaH significantly reduced virulence and impaired the pathogen’s ability to establish persistent infection. Notably, RfaH expression increased under stress conditions, such as high osmolarity and temperature, underscoring its role in bacterial adaptation. On the other hand, the absence of RfaH led to motility defects and enhanced bacterial aggregation, suggesting alterations in surface properties. Transcriptomic analysis revealed that RfaH influences a broader set of genes beyond the O-antigen biosynthesis operon, including virulence factors critical for host adaptation. Additionally, we identified a potential non-coding RNA within the 5′-UTR of the O-antigen biosynthesis operon, which may regulate urease operon. Overall, our findings establish RfaH as a key regulator of *Y. pseudotuberculosis* virulence, shedding light on the molecular mechanisms that enable bacterial survival in challenging environments.

## Introduction

The genus *Yersiniae*, in the family *Enterobacteriaceae*, consists of 11 species, of which *Yersinia pseudotuberculosis, Y. enterocolitica*, and *Y. pestis* are known to cause diseases in mammals [1–3]. *Y. pestis*, transmitted by fleas, is the causative agent of plague [4], while both *Y. pseudotuberculosis* and *Y. enterocolitica* are enteric pathogens that are typically acquired by ingestion of contaminated food or water, leading primarily to self-limiting conditions such as adenitis, enteritis, diarrhea, and ileitis [2,5,6]. High-dose *Y. pseudotuberculosis* infection in rodents, such as guinea pigs and mouse models, results in systemic infection due to translocation from the intestinal tract to the spleen and liver [7]. The pathogenicity of all three *Yersinia* species is largely dependent on the Type III Secretion System (T3SS) and its related virulence substrate proteins, known as *Yersinia* outer proteins (Yops), which are encoded on a 70-kb virulence plasmid [8,9]. Low-dose *Y. pseudotuberculosis* oral infections cause chronic infection in mouse cecal tissue without signs of disease [10]. This model of *Y. pseudotuberculosis* persistent infection offers insight into bacterial mechanisms of importance for initiation and maintenance of persistent infections. We have previously explored the nature of *Y. pseudotuberculosis* persistent infection by profiling transcriptional changes from the early to the later persistent phase of infection [11]. The study identified potentially significant key players during persistence, which primarily reflected environmental conditions the pathogen encountered in the cecal lymphoid follicles. A set of global transcriptional regulators, such as the Crp/CsrA/RovA cascade, regulated gene expression, allowing *Y. pseudotuberculosis* to adapt to long-term residence in the host. Notably, expression of the gene encoding the transcriptional regulator RfaH was significantly enhanced during the persistent state of infection, but was not investigated further [11].

RfaH is a transcriptional regulator belonging to the NusG family of proteins [12]. It enables RNA polymerase to bypass intrinsic terminator sites or DNA binding proteins, facilitating the complete transcription of long operons [13,14]. The specificity of RfaH for its target genes depends upon a 12-nucleotide conserved regulatory site called the operon polarity suppressor (*ops*), which is typically located at upstream regions of the operons regulated by RfaH [15,16]. RfaH prevents early transcriptional termination and enhances transcriptional elongation, ensuring, for example, lipopolysaccharides (LPS) biosynthesis, shown for many bacteria [17]. In addition, RfaH has been linked to various roles, including initiation of translation by interacting with the 30S ribosome [12], regulation of operons involved in capsular biosynthesis [18], hemin uptake components [19], and production of toxins like hemolysins and cytotoxic necrotizing factor [20]. Indeed, survival of *Vibrio vulnificus* in serum is dependent on RfaH [21]. RfaH has also been implicated in the pathogenesis of various pathogens where its loss leads to attenuation in virulence of *Salmonella* [19], *E. coli* [22], and *Klebsiella* [23]. In *Y. pseudotuberculosis*, RfaH has been reported to contribute to resistance against antimicrobial chemokines and survival during mouse infections [24]. However, the precise mechanisms by which RfaH enhances expression of these components remain incompletely understood. Despite significant scientific advances in understanding the RfaH mechanism in various pathogens, knowledge about its molecular mechanisms in microbial pathogenesis, particularly *Y. pseudotuberculosis*, remains limited [25]. In this study, we conducted phenotypic assays to dissect RfaH-dependent changes during bacterial growth and the establishment of infection in mouse model. Furthermore, we extended our understanding of RfaH’s role in the pathogenesis of *Y. pseudotuberculosis* through transcriptomic analysis by performing transcriptomic profiling of wt, *rfaH* deletion mutants, and various RfaH related mutant strains of *Y. pseudotuberculosios* at both environmental (26°C) and host body (37°C) temperatures to identify RfaH regulon. We identified common and mutation-specific genes and pathways, shedding light on the potential molecular mechanisms of RfaH in persistent *Y. pseudotuberculosis* infections. Our findings provide a deeper understanding of RfaH’s role in *Y. pseudotuberculosis* virulence mechanisms and offer a valuable molecular reference for future studies to understand persistent infections caused by this pathogen.

## Results

### RfaH expression is growth phase-dependent and can be induced by envelope stress

Given that the *in vivo* transcriptome profile of persistent *Y. pseudotuberculosis* resembled that of stationary phase bacteria [11], we investigated an eventual growth phase dependence of RfaH expression in *Y. pseudotuberculosis*. As a result, the expression of RfaH increased over time and peaked at the late stationary phase (Figure 1A), similar to the pattern observed for *rfaH* in *S. enterica* serovar Typhimurium [26,27]. This provided a starting point for further investigation into the environmental cues that might influence production of RfaH. Moreover, using *in vitro* settings that mimicked various host environments, we investigated the impact of specific environmental factors such as high osmolarity, bile salts, glucose, low pH, oxidative stress, temperature, and a combination of high osmolarity, bile salts, and glucose (BNG) on RfaH expression in bacterial cultures at late stationary phase. Surprisingly, all the conditions tested increased RfaH expression (Figure 1B). Thus, host stress conditions induce the expression of RfaH, indicating that this regulatory protein may play a role in the pathogen’s adaptability to hostile environments in the host.

**Figure 1.**
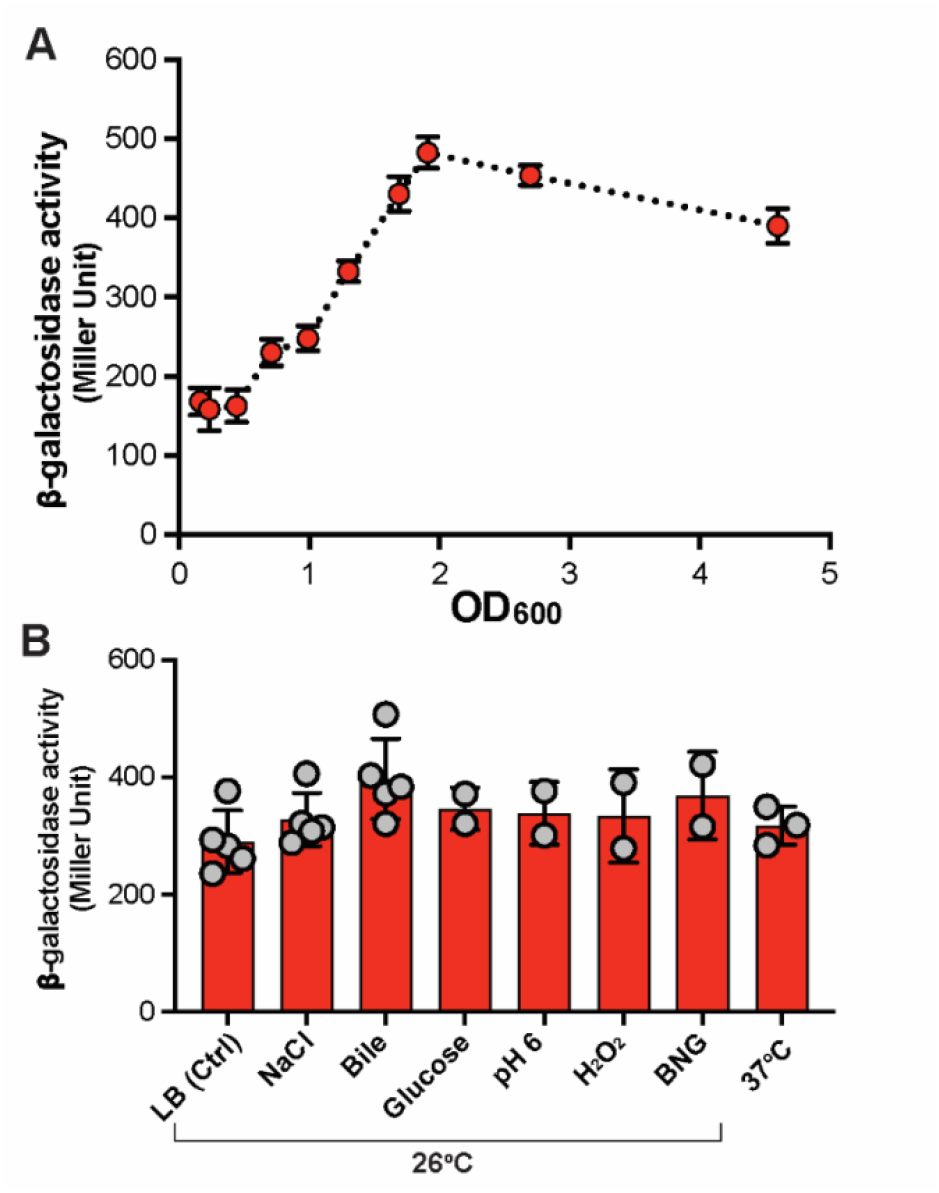
RfaH expression increased during the stationary phase and under stressful conditions. **A** The β-galactosidase activity of the *rfaH* promoter fused with promoterless lacZ reflected in the hourly increase in optical density (600_nm_) during bacterial growth at 26°C. **B** The β-galactosidase activity of the *rfaH* promoter in bacterial cultures treated with 125 mM NaCl, 0,5% bile salts, 0,4% glucose, pH:6, 10 mM H_2_O_2_, and a combination of 125 mM NaCl, 0,5% bile salts, and 0,4% glucose (BNG). The dots in A and bars in B represent mean values, and error bars show standard deviations.

### Lack of *rfaH* is associated with bacterial clustering and a non-swimming phenotype

To investigate the impact of *rfaH* deletion, we performed phenotypic assays, including light and atomic force microscopy, to assess bacterial morphology. Assays were conducted under conditions with the highest RfaH expression among those tested, and a control condition at 26°C. As a result, the *ΔrfaH* strain formed cluster-like structures after being subjected to NaCl, BNG, and temperature rise to 37°C for two hours. These findings imply that RfaH affects the surface structures of bacteria. In addition, when treated with the three environmental stresses, *ΔrfaH* lost its ability to swim in liquid media (Figure 2A). Accordingly, expression of *rfaH* in trans could complement the clustering and swimming phenotype (Figure 2A). To evaluate motility, wt and *ΔrfaH* strains were subjected to soft agar motility assays after exposure to NaCl and BNG. This was performed at 26°C to ensure motility since *Y. pseudotuberculosis* is non-motile at 37°C [11]. The *ΔrfaH* strain showed reduced motility, which in consistence with the non-swimming phenotype, was rescued by *rfaH* trans complementation (Figure 2B). To detect any potential impacts on flagella, we subjected both non-treated bacteria and bacteria treated with NaCl and BNG to atomic force microscopy. We observed that the non-swimming phenotype was not due to a lack of flagella, as the *ΔrfaH* strain had intact flagella in NaCl and BNG supplementation media (Figure 2C). Despite intact flagella, the observed reduction in *ΔrfaH* motility, suggests that RfaH may be involved in regulating surface structures crucial for efficient swimming, potentially through its influence on cell aggregation or chemotaxis signaling pathways.

**Figure 2.**
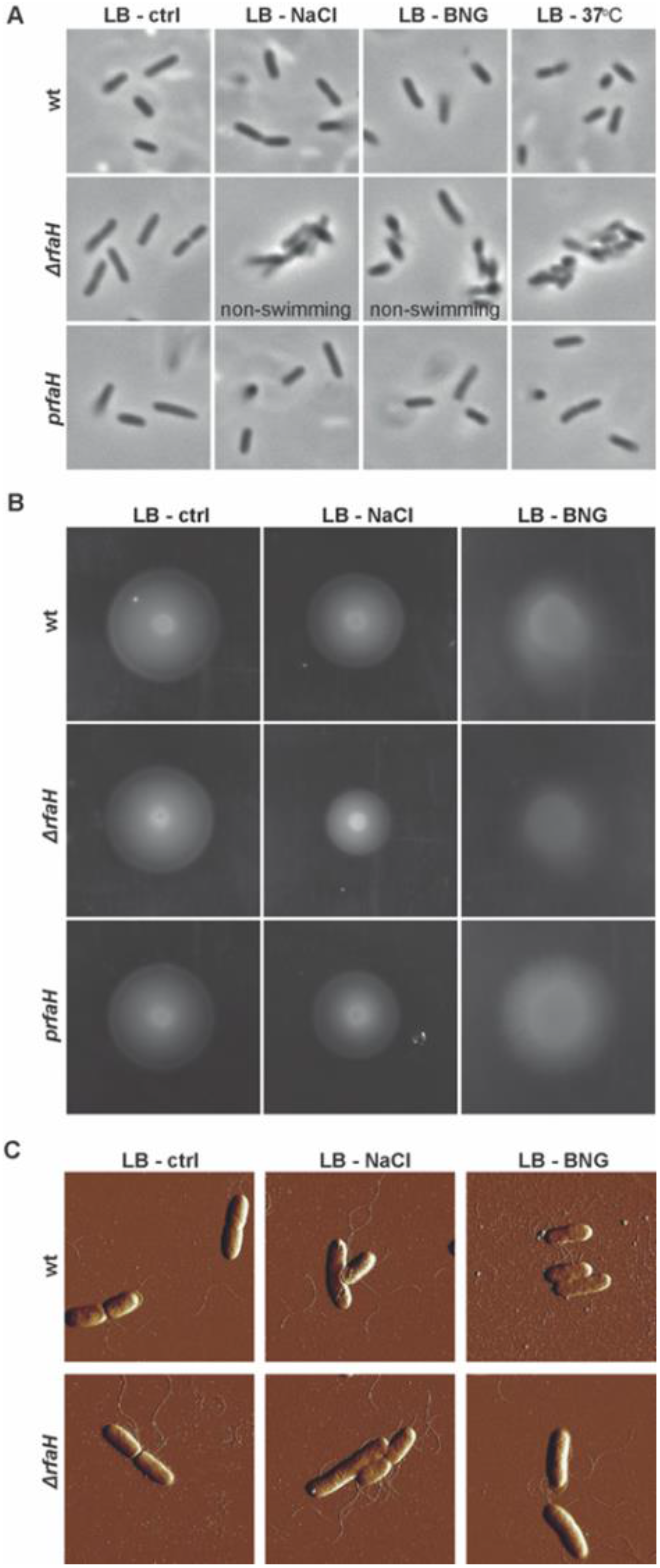
Loss of *rfaH* leads to reduced motility in the presence of intact flagella under stress conditions. **A** Phase contrast microscopy images of wt, Δ*rfaH*, and Δ*rfaH* complemented with trans *rfaH* expression *(prfaH)* strains under control conditions at 26°C, 125 mM NaCl, a combination of 125 mM NaCl, 0,5% bile salts, and 0,4% glucose (BNG) and 37°C. **B** Motility profile of wt, Δ*rfaH*, and *prfaH* under 125 mM NaCl and BNG along with control condition at 26°C. **C** The atomic force microscopy images of the flagella for the wt and Δ*rfaH* strains under 125 mM NaCl and BNG along with control conditions at 26°C.

### Defective O-antigen biosynthesis contributes to *rfaH*-dependent phenotypic changes

Given that RfaH is known to regulate LPS in *Y. pseudotuberculosis* [24], we hypothesized that the cluster formation of the *ΔrfaH* mutant strain might be due to a defect in LPS biosynthesis. LPS comprises three distinct regions: lipid A, a core polysaccharide, and an O-antigen (Figure 3A). The comparison of the LPS profiles of the wt and *ΔrfaH* strains using SDS-PAGE and silver staining revealed that the *ΔrfaH* strain produced lower molecular weight LPS molecules in comparison to the wt strain at both 26°C and 37°C (Figure 3B). Further investigation into the composition of Lipid A species in both wt and *ΔrfaH* strains using MALDI-TOF analysis revealed no differences between the two strains at either temperature (Figure 3C and D). This suggests that the lower molecular weight of the LPS produced by the *ΔrfaH* strain may result from insufficient O-antigen biosynthesis, as the operon responsible for O-antigen biosynthesis that is known to be regulated by RfaH, has an *ops* sequence located in the upstream region of the operon (Figure 3E).

**Figure 3.**
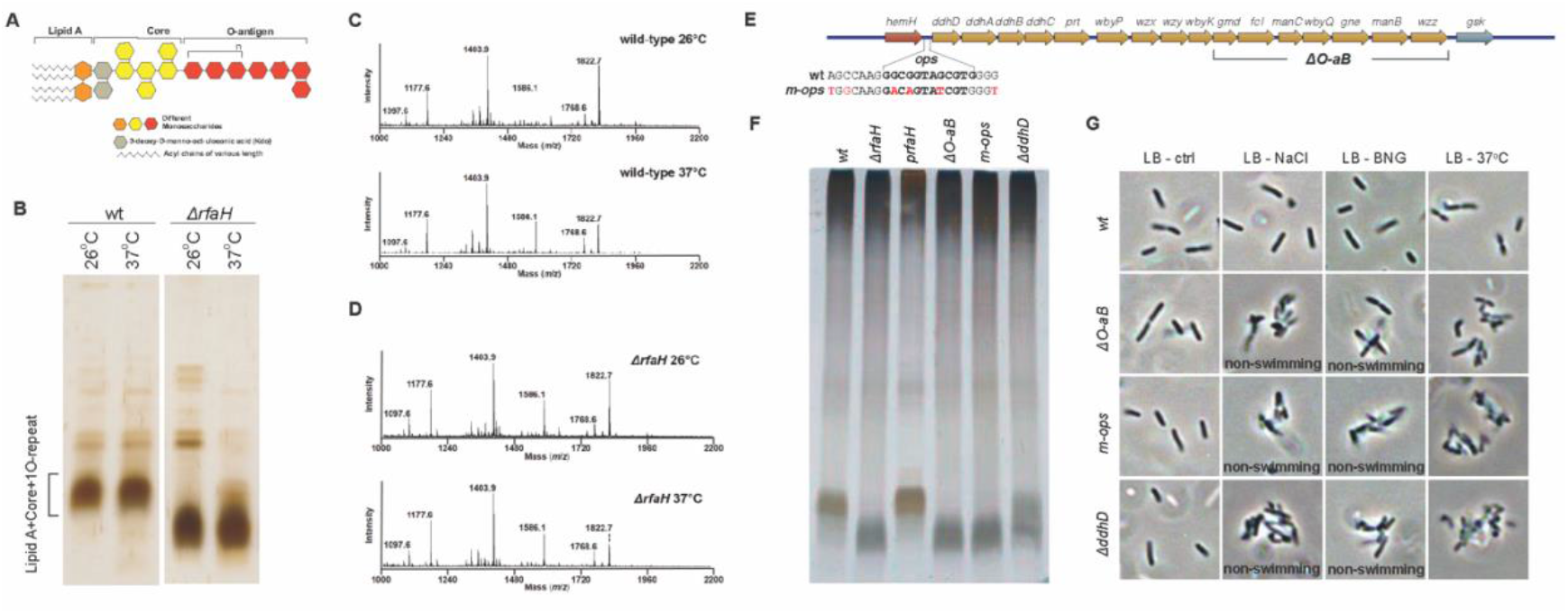
RfaH-related phenotypic changes are indirectly impacted by defective O-antigen biosynthesis. **A** Composition of LPS in *Y. pseudotuberculosis*. **B** LPS profiles of wt and *ΔrfaH* mutant strains were analyzed at 26°C and 37°C in 15% SDS-PAGE with silver staining. **C** Representative MALDI-TOF mass spectra of Lipid-A for wt and **D** for *ΔrfaH* analyzed at 26°C and 37°C. **E** Schematic presentation of O-antigen biosynthesis with the *ops* sequence located on the upstream region. The nucleotide substitutions introduced in the *ops* sequence to generate *m-ops* mutant are marked in red. The array of seven genes on the distal part of the operon is shown for the *ΔO-aB* deletion mutant. **F** LPS profiles of wt, Δ*rfaH*, and other O-antigen biosynthesis operon mutants and *ΔrfaH* complementation strain were analyzed at 26°C in 15% SDS-PAGE with silver staining. **G** Phase-contrast microscopy images of multiple mutants with defective O-antigen biosynthesis.

To determine if the observed phenotypes of *ΔrfaH* result from a defect in O-antigen biosynthesis, we set out to construct mutant strains deficient in O-antigen biosynthesis. To do this, we deleted the gene encoding for CDP-6-deoxy-delta 3,4-glucoseen reductase (*ΔddhD*), an enzyme essential to the synthesis of CDP-ascarylose during the biosynthesis of O-antigens [28]. Moreover, we deleted a sizable DNA region, including seven genes (*gmd, fcI, manC, wbyQ, gne, manB*, and *wzz*), toward the end of the O-antigen operon to severely attenuate O-antigen biosynthesis (called *ΔO-aB*) (Figure 3E). Additionally, we generated a strain with a modified *ops* sequence located upstream of the O-antigen biosynthesis operon by introducing six nucleotide substitutions inside and in flanking regions of the *op’s* sequence (called *m-ops*) (Figure 3E and 3F). LPS profiling of the mutant strains showed that all of them had deficits with lower molecular weight LPS profiles similar to that seen for *ΔrfaH* (Figure 3F). Also, the defective LPS profile of the *m-ops* strain demonstrated that RfaH positively regulates this O-antigen biosynthesis operon. Finally, microscopy analysis of the several O-antigen mutants subjected to NaCl and BNG treatments and 37°C revealed phenotypes comparable to those of the *ΔrfaH* strain. Following BNG treatment, these mutants similarly exhibited the clumping and non-swimming phenotype (Figure 3G). Thus, the clumping and non-swimming phenotypes observed for *ΔrfaH* could be due to indirect effects from defective O-antigen biosynthesis in the *ΔrfaH* strain.

### The global effect of RfaH on the *Y. pseudotuberculosis* transcriptome

To investigate other potential genes or operons regulated by RfaH, we performed RNA-seq on *ΔrfaH* and wt strains at 26°C and 37°C temperatures. Furthermore, to filter genes whose expression regulation is affected by defective O-antigen biosynthesis in the absence of RfaH, we included *m-ops* and *ΔO-aB* strains with defects in O-antigen biosynthesis. Differential expression analysis revealed reduced expression of all O-antigen biosynthesis operon genes in *ΔrfaH* and *m-ops* mutants relative to the wild-type at both 26°C and 37°C (Figure 4A). As expected, expression of only seven deleted genes of O-antigen biosynthesis operon was reduced in the ΔO-aB strain (Figure 4A). To identify genes that are exclusively differentially regulated in *ΔrfaH*, we compared differentially regulated genes (DEGs) in *ΔrfaH* to those regulated in *m-ops* and *ΔO-aB* under both 26°C and 37°C to exclude genes whose expression is affected by deficient O-antigen biosynthesis but not RfaH. This analysis revealed 8 DEGs exclusively related to *ΔrfaH* mutation at 26°C (Figure 4B). Among the 8 DEGs, six were downregulated (including thiosulfate transporter, cytoplasmic protein, and hemolysin), while genes encoding transposase and lipoprotein were upregulated (Table S1). On the other hand, we identified 23 DEGs exclusively associated with *ΔrfaH* mutation at 37°C (Figure 4C), indicating a potentially broader regulatory role for RfaH at host body temperature. Of the 23 DEGs, 10 were downregulated (including tail E family protein, monosaccharide-transporting ATPase, and holin), and 13 were upregulated (including type 11 methyltransferase, phage transcriptional regulator AlpA, type VI secretion protein and fimbrial protein) (Table S1).

**Figure 4.**
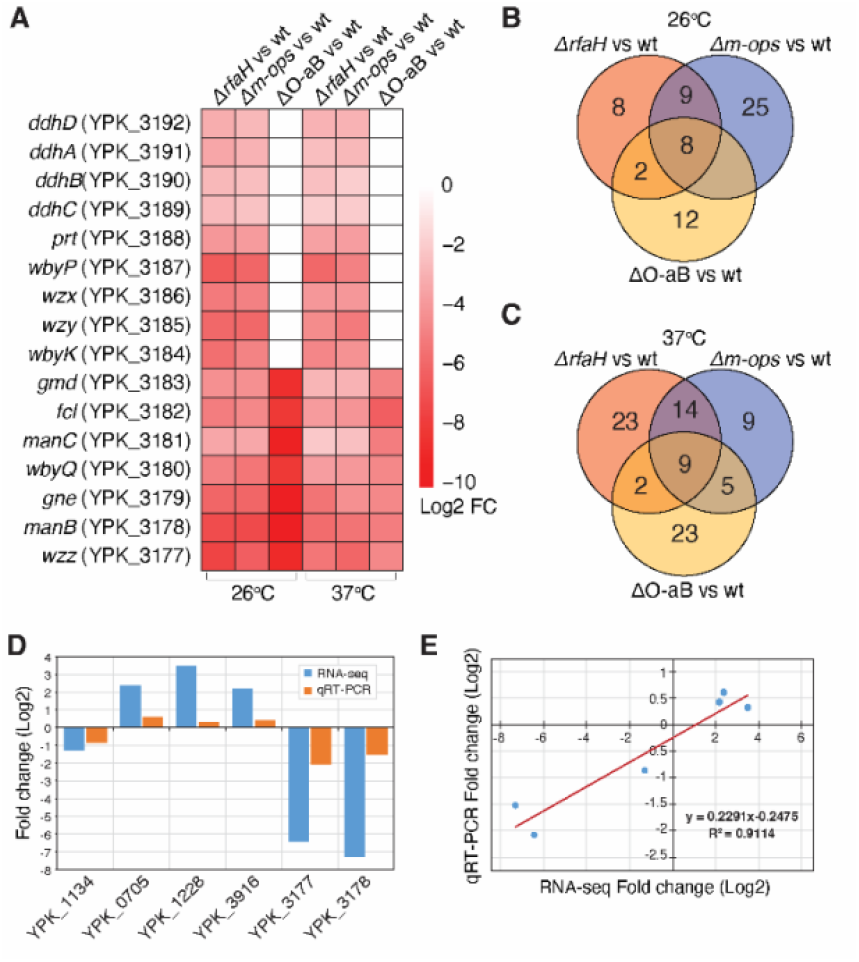
Differential gene expression profile of *ΔrfaH, m-ops, ΔO-aB* mutants, and wild-type strains with validation of the gene expression results by qRT-PCR. **A** Heatmap highlighting low expression profiles of O-antigen biosynthesis operon genes in *ΔrfaH, m-ops*, and *ΔO-aB* versus the wt strain at 26°C and 37°C. **B** Venn diagram highlighting gene expression patterns of *ΔrfaH, m-ops*, and *ΔO-aB* versus the wt strain at 26°C and **C** at 37°C. **D** Bar plot of RNA-seq and qRT-PCR expression levels. The X-axis represents seven randomly selected genes, and the Y-axis represents the log2 foldchange from RNA-seq and qRT-PCR. **E** The linear regression analysis of expression level between RNA-seq and qRT-PCR data. The X-axis represents the log2 fold change of RNA-seq, and the Y-axis indicates the log2 fold change of qRT-PCR. Abbreviations: vs-versus, wt-wild type.

Interestingly, the same analysis identified 25 DEGs exclusively related to *m-ops* strain at 26°C (Figure 4B). They are unlikely to be affected by deficient O-antigen biosynthesis since we filtered out DEGs commonly regulated in *m-ops, ΔrfaH*, or *ΔO-aB* mutants. This excludes the effects of impaired O-antigen biosynthesis and suggests a potential additional regulatory role for the hypothetically expressed wt *ops* sequence. Among the 25 DEGs, most of the *ure* operon [29] genes were downregulated, including *ureC, ureE, ureF, ureG*, and *ureD*. Notably, only nine DEGs were exclusively related to *m-ops* strain at 37°C (Figure 4B and C, and Table S2). Similarly, we identified 35 DEGs exclusively related to *ΔO-aB* mutation at both temperatures (Figure 4B and C, and Table S3). The relatively higher number of DEGs in the *ΔO-aB* strain could the consequence of the complete lack of O-antigen biosynthesis, as this strain lacks seven genes encoding proteins critical for O-antigen biosynthesis compared to *ΔrfaH* and *m-ops* with less O-antigen biosynthesis. Eight DEGs were common across all three mutant strains compared to the wt strain at 26°C and 37°C (Figure 4B and C). All these were related to the O antigen biosynthetic process and downregulated in all the strains (Table S4). To validate the accuracy and reliability of the gene expression inferred from our RNA-seq data, seven genes were randomly selected and analyzed by qRT-PCR. The expression pattern revealed by qRT-PCR analysis was consistent with RNA-seq (Figure 4D). In addition, the significant correlation observed (R^2^ = 0.8807) between RNA-seq and qRT-PCR data (Figure 4E) indicates the robustness and reliability of the DEGs identified in this study using RNA-seq.

### *rfaH* is required for full *in vivo* virulence of *Y. pseudotuberculosis*

To test the importance of RfaH in *Y. pseudotuberculosis* virulence in a mouse model, we performed low-dose oral infection in FVB/n mice with *ΔrfaH* mutant strain. Thereafter, we followed *Y. pseudotuberculosis* colonization with the presence or absence of a bioluminescent signal produced by a bioluminescent reporter system introduced in virulence plasmid and monitored with In Vivo Imaging System (IVIS). We observed that only 19% of the mice were initially infected with the *ΔrfaH* mutant strain, compared to 82% of the ones infected with the wt strain (Figure 5A, Figure S1A). On the other hand, 36% of wt-infected mice developed acute infection by 16 days post-infection (dpi) and succumbed to the infection, while only one of the *ΔrfaH* infected mice showed symptoms of acute infection (Figure 5A, Figure S1A). The ability of *ΔrfaH* strain to establish persistent infection diminished by less than half as only two mice infected with the *ΔrfaH* strain could keep the infection up to 30 dpi, while five mice infected with the wt strain developed persistent infection (Figure 5A, Figure S1A). This reduction is likely influenced by the requirement for *rfaH* during the initial stages of infection, with a notable number of mice remaining uninfected. Moreover, we observed that the attenuation of *ΔrfaH* in the mouse oral infection model could be complemented with the expression of *rfaH in trans* (Figure 5A, Figure S1A). As a result of complementation, four mice developed persistent infection, indicating that RfaH is required to establish persistent infection in mice. Additionally, we monitored the presence of *prfaH* during infection by continually screening for antibiotic resistance, encoded on the plasmid that encodes *rfaH*, on *Y. pseudotuberculosis* that was shed through feces (Figure S1B). Our results confirmed that *prfaH* was still present even after 31 dpi and with a high bacterial load after 10 dpi (Figure S1B). Further analysis of the survival of mice infected with different strains showed significantly higher survival of mice infected with *ΔrfaH* compared to mice infected with strains expressing this regulator (Figure 5B).

**Figure 5.**
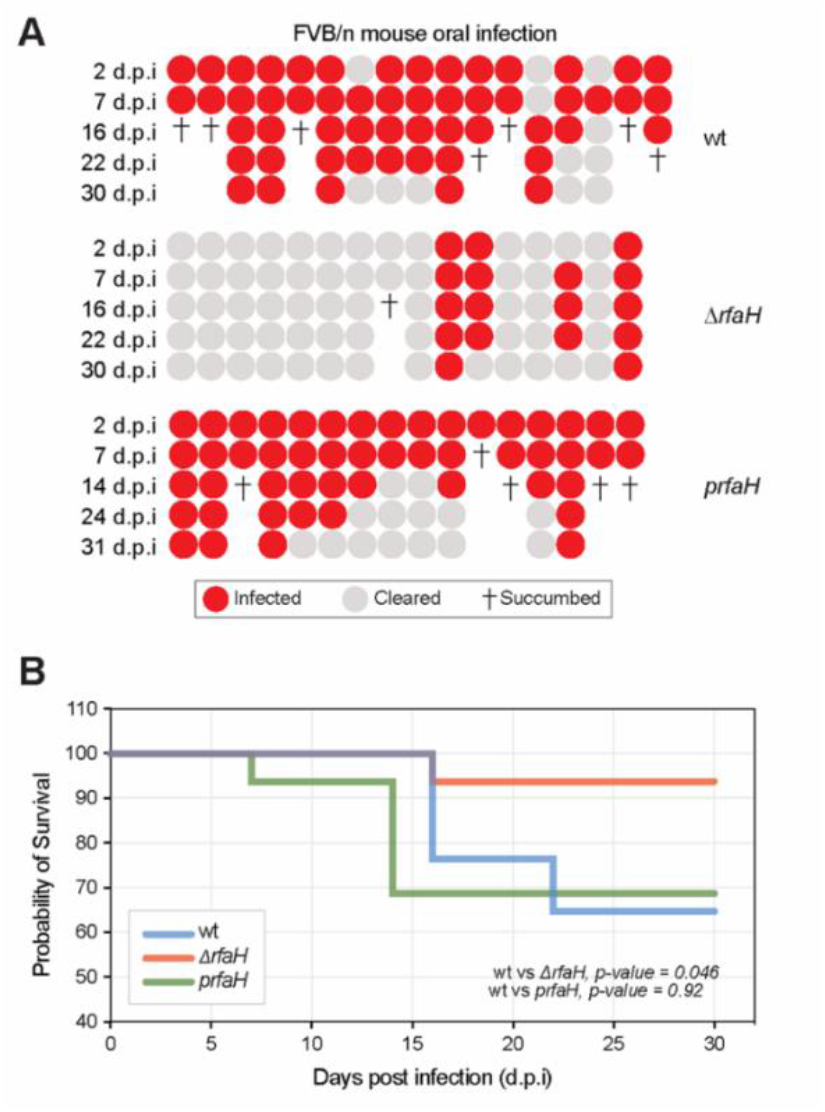
The lack of RfaH leads to virulence attenuation in low-dose oral infection of FVB/n mice and can be complemented by trans expression of rfaH. **A** Real-time IVIS monitoring of low-dose oral infection of *Y. pseudotuberculosis*. Mice were infected orally with overnight bacteria cultures, and the infection process was monitored every day post-infection in wt, *rfaH* mutant, and *prfaH* strains. Each circle represents an individual mouse infected in either wt (17), rfaH mutant (16), or prfaH (16). The red and grey colors indicate positive and negative bioluminescent signals detected by IVIS, respectively. Each cross symbol indicates an individual mouse showing infection symptoms such as fussy fur, diarrhea, and weight loss and therefore, succumbing to the infection. **B** Survival percentages of mice infected with wt, *ΔrfaH*, and *prfaH* strains. The Log-rank (Mantel-Cox) test was used to calculate the *p*-value.

### 5’-UTR of O-antigen biosynthesis operon encodes a hypothetical small RNA

Mutations in the *ops* region can affect the binding of RfaH, leading to premature transcription termination or improper transcription of downstream genes, thereby reducing their expression. Interestingly, our RNAseq data analysis revealed that the *ops* mutation affected the transcriptional regulation of several genes that are not regulated by RfaH, including downregulation of the urease operons (*ureC, ureE, ureF, ureG*, and *ureD*), citrate reductase cytochrome c, glycerol kinase, and glycosidase in the *m-ops* strain compared to the wt at 26°C (Table S2). This finding underscores a distinct regulatory role for this *ops* element, independent of RfaH, in expression of specific genes. Notably, analysis of read mappings surrounding the *ops* region revealed a unique mapping profile in the *m-ops* strain. The relatively low coverage of the GGGGGG sequence downstream of the *ops* sequence in the wt and *ΔrfaH* strains, compared to the flanking regions, was absent in the *m-ops* strain (Figure 6). This observation suggests the presence of a potential RNA motif that may have been disrupted by a single nucleotide substitution in the *m-ops* strain, possibly indicating the action of an RNase that processes the 5’ UTR of the O-antigen biosynthesis operon. Consequently, the cleavage of this region, resulting in an approximately 136 nt long 5’ UTR, may function as a regulatory non-coding RNA precisely modulating the expression of genes uniquely regulated in the *m-ops* strain. Although this hypothesis is beyond the scope of the current study, it requires further investigation to elucidate the novel regulatory role of the 5’ UTR in the O-antigen biosynthesis operon.

**Figure 6.**
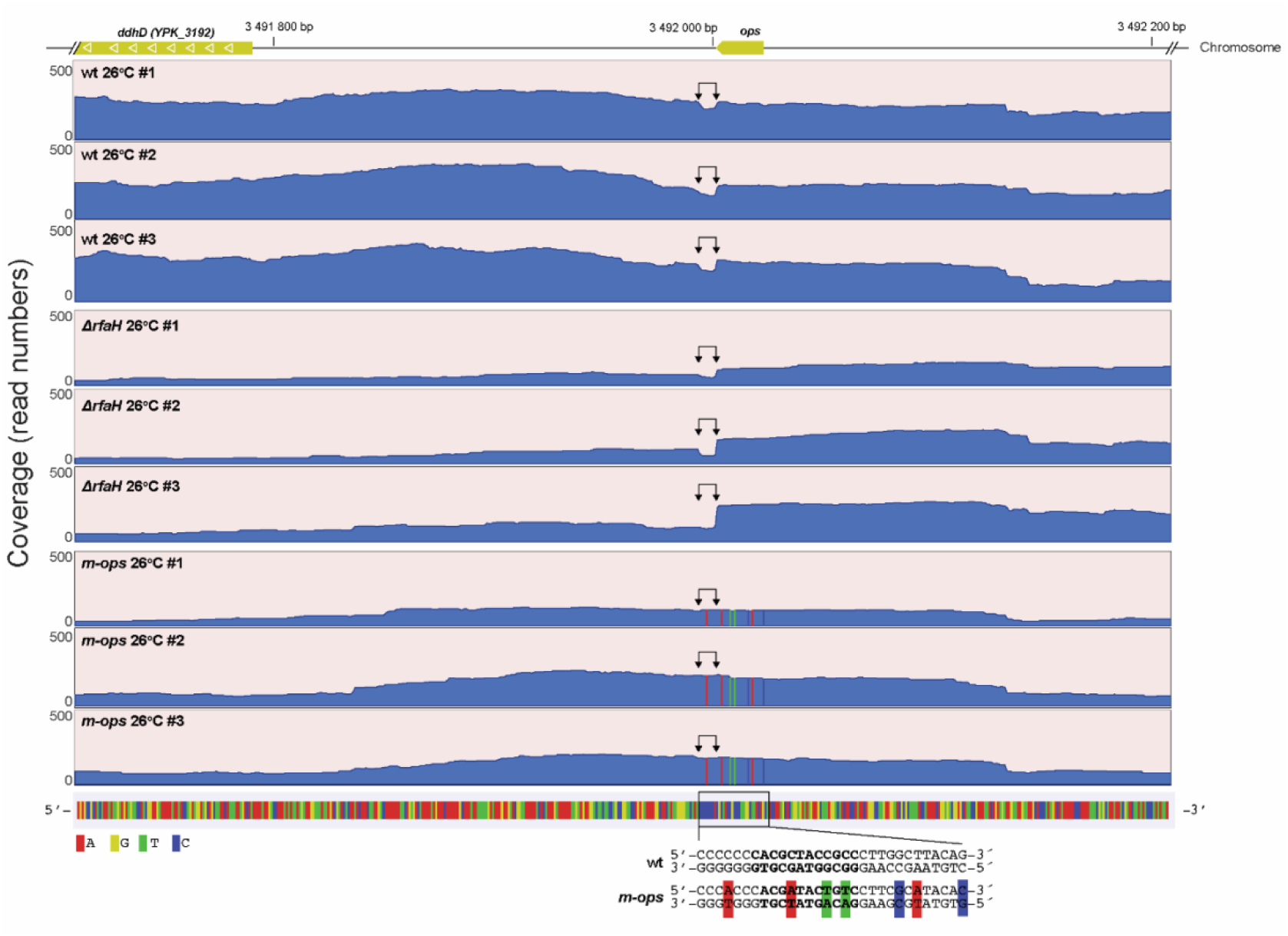
RNA-seq read mappings of the O-antigen biosynthesis operon’s 5’ UTR indicate the presence of a hypothetical non-coding RNA. Read coverage tracks (blue) for the 5’ UTR of the O-antigen biosynthesis operon (upstream of *ddhD*) and the *ops* sequence across triplicates of wt, *ΔrfaH*, and *m-ops* strains at 26°C. The relatively low coverage of the GGGGGGGG motif downstream of the *ops* sequence in the wt and *ΔrfaH* strains is highlighted by two arrowheads in each track. In contrast, this low coverage of the GGGGGGG sequence is absent in the *m-ops* strain, likely due to a point mutation that results in the GGGTGGG sequence. The bottom track displays the DNA sequence with nucleotides represented in distinct colors. The *ops* sequence and its flanking regions in the wt and *m-ops* strains are delineated below, with the *ops* sequence in bold and nucleotide substitutions in the *m-ops* strain indicated by color coding.

## Discussion

Previous studies have reported that RfaH is a key regulator of several operons, whose expression is fully dependent on the presence of this regulatory protein [25]. Our previous *in vivo* transcriptome analysis revealed that the gene encoding the transcriptional antiterminator RfaH was upregulated during persistent *Y. pseudotuberculosis* infection [11], suggesting that this gene might play an essential role in perpetuating the infection. Here, we showed that the expression of RfaH reaches the highest level at the stationary phase when comparing different infection related *in vitro* conditions. This correlates with the observed upregulation of RfaH during persistent infection, where the transcriptome of persistently colonized *Y. pseudotuberculosis* is more similar to bacteria in the stationary phase than to that in growing bacteria [11]. Similarly, Rojas et al. [26] reported that the expression of RfaH increased up to two-fold when the growth of *Salmonella* reached the late exponential phase and remained high throughout the stationary phase. Moreover, we observed RfaH upregulation also under other stress conditions, including NaCl, bile salts, temperature, and BNG, implying a possible role of RfaH in adaptation to and maintenance in harsh conditions.

Another signature of persistent bacteria was upregulation of genes involved in envelope perturbations [11]. In this study, LPS profiling of strains mutated in different RfaH dependent features highlighted that all had defective LPS profiles, as observed for *ΔrfaH*. Moreover, the defective LPS profile of the *m-ops* strain demonstrated that RfaH positively regulates the O-antigen biosynthesis operon. Defective LPS biosynthesis impacted multiple bacterial surface processes, including clumping phenotypes of the various mutant strains, which were comparable to the *ΔrfaH* strains. In *Salmonella*, LPS regulated by the RfaH gene was reported to play a key role in the interaction between cells and the environment, contributing to virulence [26]. Similarly, regulation of LPS has been reported as a common way for bacteria to change their surface and adapt to new environments, such as increased temperature and exposure to different chemicals [30]. These results suggest that LPS-defective bacteria will have challenges to adapt during infection. Previous studies in *E. coli* have shown intestinal colonization was impacted by the regulatory action of RfaH on LPS-core production [31]. Surprisingly, the LPS defect might also be responsible for the non-swimming phenotype of *ΔrfaH*, as shown in this study. The reason for this is unclear, but it might be clustering *per se* or that the LPS defect affects the organization of membrane components in a way that hinders flagella function.

Previous studies have reported that deletion of the *rfaH* gene attenuates bacterial virulence in several ways, including reduction of capsules and intact LPS, hemin receptor, and alpha-hemolysin during infection [25]. The *E. coli* A192PP*ΔrfaH* mutant was highly susceptible, with no colonies detected after 30 minutes of incubation in 22% normal human serum or heat-inactivated serum (56°C for 30 minutes)[32]. In this study, our transcriptome analysis highlighted that deletion of *rfaH* exclusively affected the expression of a specific set of genes at 26 °C and 37°C compared to the wt strain (Table S1). These genes might shed light on the global regulatory function of RfaH in *Y. pseudotuberculosis*. Besides the O-antigen biosynthesis operon, genes encoding cytoplasmic protein, flagellar basal body rod protein *FlgF*, and hemolysin were downregulated in the *ΔrfaH* strain compared to the wt strain at 26°C. Similarly, genes encoding holin and monosaccharide-transporting ATPase were downregulated at 37°C (Table S1). In *E. coli*, the alpha-hemolysin expression has been reported to be under the control of RfaH [33]. A study on *Y. enterocolitica* reported a decrease in the amount of Mg^2+^ transport ATPase proteins and phosphate ABC transporter in the YeO3-*ΔrfaH* strain [34]. As suggested by Nagy et al.[19], this might be an adjustment in response to decreased LPS production due to the RfaH deletion, resulting in reduced demand for sugars and energy. In the *Y. pseudotuberculosis DrfaH* strain, the expression of genes encoding lipoprotein and transposase was enhanced at 26°C while those encoding type VI secretion protein, transcriptional regulator AlpA, type 11 methyltransferase and fimbrial protein were upregulated at 37°C (Table S1). A previous study of *A. baumannii* reported that a LPS-deficient strain displayed increased expression of genes encoding cell envelope and membrane biogenesis like lipoproteins and lipoprotein transport system [35]. It is therefore possible that alteration of the lipoprotein content of the outer membrane is a response to LPS loss. Fimbrial proteins and type II methyltransferases (MTases) have been reported to play key roles in the virulence and defense of several bacteria, including *E. coli* [36], *Salmonella* [37], *Avibacterium paragallinarum* [38], *Metamycoplasma hominis* [39] and *Xanthomonas axonopodis pv. Glycines* [40]. Similarly, an AlpA positively regulated a programmed cell death pathway linked to the virulence of *Pseudomonas aeruginosa* [41] while Type VI Secretion System (T6SS) have been identified within pathogenicity islands of *Salmonella* [42] as well as in survival of *Vibrio anguillarum* and *Vibrio cholerae* in stress conditions[43],[44]. Interestingly, the high number of hypothetical genes among those regulated by RfaH suggests that much remains unknown about this transcriptional regulator. However, further analysis of these genes could provide valuable insights into potential mechanisms of *rfaH* regulation in infection.

The loss of RfaH severely attenuated the virulence of *Y. pseudotuberculosis* upon infection, as only two mice infected with the *ΔrfaH* strain could keep the infection up to 30 dpi as compared to five mice infected with the wt strain (Figure 5, Figure S1A). These highlight the significance of RfaH in the initial phase of infection and in the adaptation and maintenance of the pathogen at later stages. Accordingly, the inactivation of RfaH dramatically decreased the virulence of the uropathogenic *E. coli* strain [25]. However, a previous study found that deleting the *rfaH* gene in *Y. pseudotuberculosis* did not reduce its virulence in BALB/c mice during acute infection [24]. However, that study used a different *Y. pseudotuberculosis* strain (IP32953) and a different mouse model (BALB/c) for infection. We have previously demonstrated that FVB/N mice are more suitable for establishment of persistent infection by the YPIII strain than BALB/c mice. One difference between the mouse strains used in the two studies is their major histocompatibility complex class I haplotype: BALB/c mice possess H-2^d^, while FVB/N mice have H-2^q^. The susceptibility of FVB/N mice to persistent murine encephalomyelitis virus infection and the increased resistance conferred by an H-2^b^ transgene highlight the significant role of MHC haplotype variation in determining host susceptibility to infectious agents [45]. This suggests different outcomes of *rfaH* deletions could simply be due to different mouse models used in different infections. Moreover, although *Y. pseudotuberculosis* YPIII and IP32953 strains have the same O-antigen biosynthesis operon structure and upstream *ops* sequence, IP32953 is known to be more virulent and exhibit greater colonization capacity in mice than YPIII [46]. This suggests that strain-specific factors, beyond *rfaH* and O-antigen production, significantly influence virulence in mouse models of *Y. pseudotuberculosis* infection.

## Materials and Methods

### Ethics statement

Mice were housed under the Swedish National Board for Laboratory Animals guidelines. All animal procedures were approved by the Animal Ethics Committee of Umeå University (Dnr A108-10). Mice were allowed to acclimate to the new environment for one week before the experiments.

### Strains and growth conditions

The wt strain in this study was the Kanamycin-resistant bioluminescent *Y. pseudotuberculosis* YPIII/pIBX strain (Table S5). The YPIII strain is a well-studied model for *Y. pseudotuberculosis* pathogenesis [10,11]. The strains were cultured overnight at 26°C and 220 rpm in Luria broth (LB) supplemented with 50 μg/ml kanamycin. The samples were co-cultured in the morning to a starting OD of 0.05 and grown in LB at 26°C until exponential phase before being aliquoted for several in vitro growth and total RNA purification experiments at 26 and 37°C. For stress conditions, the LB was supplemented with 125 mM NaCl, 0.5% bile, 0.2% glucose, and a combination of all three (LB-BNG).

### Mutant and other strain constructions

To generate in-frame deletion of a single gene or large genomic region of DNA, 200 nucleotides flanking region in both ends of the region of interest were PCR amplified and ligated together with *Sal*I and *Bgl*I (New England Biolabs, Inc) into pDM4 [47] by In-Fusion HD cloning Kit (Clontech Laboratories, Inc) according to manufacturer’s instructions. The same strategy was employed for multiple nucleotide substitutions by using degenerate PCR primers at the PCR amplification step. The plasmids were transformed into *E. coli* DH5αλpir and positive clones were selected on Chloramphenicol (50 μg/ml) containing agar plates, confirmed by colony PCR. Confirmed plasmid constructs were transformed in *E. coli* S17-1λpir conjugation strain for conjugal mating with *Y. pseudotuberculosis* YPIII/pIBX. Positive allelic exchange was selected as described previously [47]. The mutant strains of *ΔrfaH, m-ops*, and *ΔO-aB* were finally confirmed with sequencing (Table S5). For the complementation of the *rfaH*, the gene and its promoter sequence, were cloned into *BamH*I and *HindIII* sites of pMMB66HE resulting in *prfaH*. The promoter region of *rfaH* and a part of the downstream region encoding the first 13 amino acids were fused to promoterless *lacZ* in pFU61, where ColE1 had been exchanged with the low-copy replicon pSC101.

### Mouse infection and bioluminescent imaging

Bacteria were sub-cultured on LB agar plates supplemented with kanamycin (50 μg/ml). For infection, the bacteria were grown overnight in LB at 26°C, and concentrations were estimated by absorbance at OD_600_nm. Cultures were re-suspended to 10^7^ CFUs/ml in sterilized tap water supplemented with 150 mM NaCl. Viable count and drinking volume determined the infection dose. Eight weeks old female FVB/N (Taconic Farms, Inc) mice were deprived of food and water for 16 hours before oral infection with ~10^7^ CFUs of wt *Y. pseudotuberculosis* YPIII (pIBX, harboring the luxCDBAE operon in the virulence plasmid pYV) or the corresponding *ΔrfaH* mutant strain, which were supplied in their drinking water for 6 hours. Mice were inspected frequently for signs of infection and to ensure that infected mice showing prominent clinical signs were euthanized promptly to prevent suffering. The infections were monitored up to 31 days post-infection (dpi) by an *in vivo* imaging system, IVIS Spectrum, which was based on detecting the light produced by luciferase activity encoded on the *luxCDBAE* operon [48]. Before imaging, the mice were anaesthetized using the XGI-8 gas anesthesia system (Caliper LifeSciences, Inc.), which allowed control throughout anesthesia. The oxygen mixed with 2.5% IsoFloVet (Orion Pharma, Abbott Laboratories Ltd, Great Britain) was used for the initial anesthesia, and 0.5% isoflurane in oxygen was used during imaging.

### β-galactosidase assays

The *in vivo* transcriptome profile of persistent *Y. pseudotuberculosis* has been reported to resemble that of stationary-phase bacteria [11]. Thus, we created a reporter vector with the *rfaH* promoter linked to the *lacZ* gene to screen for induction of *rfaH* expression by measuring β-galactosidase activity. The reporter vector was expressed *in trans* in wt *Y. pseudotuberculosis*, and the β-galactosidase activity was measured during growth in Luria Broth (LB) at 26°C. The expression level of *rfaH* was calculated by measuring the β-galactosidase activity of the strain with the *lacZ* gene fused to the *rfaH* promoter. The β-galactosidase activity in this study was measured according to the method described by Zhang et al.[49]. β-galactosidase activity samples were taken every hour for growth phase-dependent and absorbance at OD_600_ nm was recorded. On the other hand, we measured the *rfaH* promoter activity after adding 125 mM NaCl, 0,5% Bile salts, temperature (37°C), and a combination of Bile, NaCl, and Glucose (BNG). The treatments were performed on bacterial cultures in LB with an OD_600_ of 0.8 and lasted for two hours, whereafter the activity of β-galactosidase was measured. All the experiments were repeated thrice, and enzyme activities (Miller units), were normalized according to cell density (OD_600_).

### Phenotypic analysis

The wt and *ΔrfaH Y. pseudotuberculosis* strains were grown as described in the strain and growth conditions section. To visualize bacterial morphologies, the bacteria samples were mounted on slides and visualized under a phase contrast microscope after two hours of exposure to stress conditions and when the bacterial culture reached OD_600_ = 0.8.

### Motility assay

Bacteria from overnight cultures were inoculated into LB and grown to OD_600_ = 0.8. Subsequently, they were treated with and without NaCl or BNG at 26°C for two hours. A 5 μl aliquot of each culture was spotted on LB with 0.25% agar. Plates were incubated at 26°C under aerobic conditions for 48 hours. The images of bacteria on the plates were monitored using a ChemiDoc XRS System (Bio-Rad).

### Visualization of flagella atomic force microscopy

The wt and *ΔrfaH* of *Y. pseudotuberculosis* YPIII-pIBX strains were grown overnight, diluted 25 times with LB media in the morning, and then cultured for two hours at 26°C to an OD_600_ = 0.8. Immediately after the bacterial cultures reached the desired OD_600_ value, the growth medium was supplemented with 125 mM NaCl or 0.5% bile, 125 mM NaCl, and 0.2% glucose. Following a two-hour treatment, the samples were centrifuged for four minutes at 1500 rpm, washed once with 2 mM MgCl_2_, and re-suspended in 50-200 μl of the same solution. Ten microliters of each sample were placed on freshly cleaved ruby red mica (Goodfellow Cambridge Ltd, Cambridge), incubated for 5 min at room temperature, and blotted dry before being placed into a desiccator for at least two hours. Images were collected by a Nanoscope V atomic force microscopy (AFM) (Bruker software) using ScanAsyst in the air with ScanAsyst cantilevers at a scan rate of approximately 0.9-1 Hz. The resulting images were flattened and/or plane-fitted in both axes using Bruker software and presented in amplitude (error) mode.

### LPS analysis

Diluted overnight cultures were grown at 26°C until OD_600_ = 0.8. Thereafter, 1 ml cultures were centrifuged, and pellets were re-suspended in 100 μl lysis buffer (100 mM Tris-HCl pH:6.8, 1.5% SDS, 1.5% β-mercaptoethanol) and boiled for 10 minutes. After that, samples were subjected to an overnight treatment with 80 μg Proteinase K followed by SDS-PAGE gel electrophoresis. The gels were stained with Pierce Silver Staining Kit (Life Technologies, Inc.) according to the manufacturer’s instructions.

### MALDIxin test

The bacterial pellet was resuspended in 200 μL of distilled water, washed three times with double-distilled water, and re-suspended in 100 μL of double-distilled water. A 50 μL aliquot was then submitted to mild-acid hydrolysis by adding 50 μL of 2% acetic acid in double-distilled water and heating for 1 hour at 100°C. The hydrolyzed cells were spun, the supernatant was discarded, and the pellet was suspended in 25 μL of double-distilled water. An aliquot of 0.4 μL of the bacterial solution was loaded onto the target and immediately overlaid with 0.8 μL of a super-2,5-dihydroxybenzoic acid matrix (Sigma–Aldrich, Gillingham, UK) used at a final concentration of 10 mg/mL in chloroform/methanol (90:10, v/v). The bacterial solution and matrix were mixed directly on the target by pipetting and the mix was dried gently under a stream of air (<1 min). Using the reflectron mode, MALDI-TOF MS analysis was performed on a 4800 Proteomics Analyzer (Applied Biosystems, Foster City, CA, USA). Samples were analyzed by operating at 20 kV in the negative-ion mode using an extraction delay set at 20 ns. MS data were analyzed using Data Explorer version 4.9 (Applied Biosystems)

### RNA Extraction, Library Construction, and Illumina Sequencing

Total bacterial RNAs for RNA-seq and qRT-PCR analysis were isolated from bacterial cultures grown as described above. Three biological replicates were used to represent each sample at 26°C (control) and 37°C resulting in 24 samples in total and 6 pairwise comparisons. The cultures were homogenized with 0.1 mm-sized glass beads with a Mini-Beadbeater (Biospec Products, Inc). TRIzol Reagent (Life Technologies, Inc.) was used to isolate total RNA according to the manufacturer’s instructions, followed by DNase I treatment (Roche). The NanoDrop 1000 (NanoDrop Technologies, Wilmington, DE, USA) and Agilent 2100 Bioanalyzer (Agilent Technologies, Santa Clara, CA, USA) were used to assess RNA concentration and integrity, respectively of the 24 samples. In addition, the integrity of RNA samples was further tested in agarose gel before being used for subsequent analysis.

For the library preparation, the bacterial mRNAs were enriched by depleting ribosomal RNA using a Ribo-Zero™ rRNA removal kit (Illumina). Following depletion, 2.5 μg of total RNA from each sample in duplicate was utilized as the initial material for creating cDNA libraries. The strand-specific cDNA libraries of *Y. pseudotuberculosis* were constructed using the ScriptSeq Complete Bacteria Kit (Illumina) according to the manufacturer’s instructions, while the library concentrations were measured with a Qubit 2.0 Fluorometer (Life Technologies, Inc). Finally, 10 pmol libraries were used within Illumina Reagent Kit v3 (150 cycles) and sequenced with the MiSeq System (Illumina, Inc.), and adapters were trimmed by MiSeq internal software.

### Reads Processing, Mapping, and Gene Expression Quantification

In this study, the reference genome of *Y. pseudotuberculosis* YPIII (NC_010465 for chromosome and NC_006153 for pYV plasmid) was used. The CLC-Bio Genomic Workbench (QIAGEN) was then utilized for quality, ambiguity, length trimming, alignment with the reference genome, and normalization of reads per kilobase per million mapped reads (RPKM). The CLC-Bio Genomic Workbench was used with default parameters for quality, ambiguity, and length trimming. The rRNAs and tRNAs annotations were deleted from the genome annotation files before mapping to avoid any bias related to rRNA depletion during library preparation. The Q20 percentages were more than 95%, while the Q30 base percentage, which is an indicator of the overall reproducibility and quality of the assay, was greater than 90%. Moreover, the GC contents of all the reads were above 45% of all 24 samples. Differential expression analysis was performed using the DESeq2 R package to identify DEGs between the controls and treated samples. To estimate the expression level, the DESeq2 program was used to normalize the number of counts of each sample gene using the base means. The difference was calculated, and the statistical significance was determined using the negative binomial distribution test[50,51]. Genes with a standard foldchange of less than or equal to 1 (≥1 or ≤−1) and a *p*-value of ≤ 0.05 between control and treated samples were considered differentially expressed.

### Gene Ontology (GO) Enrichment and KEGG Pathway Enrichment Analyses

The GO enrichment analysis of DEGs was conducted by integrating cluster Profiler [52]. Significantly enriched GO terms were determined by the *p*-value ≤ 0.05 with the Fisher’s exact test and the Bonferroni multi-test adjustment. Significantly enriched GO terms were assigned to the GO categories of biological process (BP), molecular function (MF), and cellular component (CC). The KEGG database was used to analyze the functional involvement of DEGs in various metabolic pathways. Furthermore, the statistical enrichment of DEGs in KEGG pathways was tested by integrating cluster Profiler [52], while the criteria for substantially enriched KEGG pathways was a *p*-value ≤ 0.05.

### Validation of DEGs by Quantitative Real-Time PCR (qRT-PCR)

To validate the reliability and repeatability of the RNA-Seq data, seven DEGs were randomly selected for verification by qRT-PCR. The bacterial strains were grown, and total RNA extracted as previously described were used as templates for cDNA synthesis with the Revert Aid H Minus First Strand cDNA Synthesis Kit (Fermentas). The gene-specific primers (Table S6) were designed using Primer Premier 5.0 software (Premier Biosoft International, Palo Alto, CA, USA). The qRT-PCR reactions were performed in triplicate for each condition using KAPA SYBR FAST qPCR Master Mix (KAPA Biosystems) and a Real-Time PCR Detection System (Bio-Rad). The stable reference genes YPK_0340 (*rpoB*) were selected as internal controls to normalize the expression data. The relative expression levels of the five DEGs were calculated according to the 2^−ΔΔCT^ (cycle threshold) method.

## Data availability

The resulting RNA-seq data files described in this study have been deposited in the Gene Expression Omnibus (GEO) database, at NCBI, under the accession number GSE272323.

## Supporting Information

**Figure S1**. The low dose oral infection of FVB/n mouse leads to virulence attenuation in *ΔrfaH* and can be complemented with in trans expression of *rfaH*. (A) Mice were infected orally with overnight cultures of bacteria and the process of infection was monitored by days post-infection (d.p.i) by detecting total photon emission, using the IVIS Spectrum system. The intensity of bioluminescent emission is represented as pseudocolors with variations in color representing light intensity; red represents the most intense light emission, while blue corresponds to the weakest signal. (B) The screening of antibiotic resistance through monitoring the presence of *prfaH* during infection. The presence of *prfaH* was constantly monitored by screening for antibiotic resistance encoded on plasmids encoding rfaH as Y. pseudotuberculosis was shed through feces.

**Table S1**. DEGs exclusively regulated in the *ΔrfaH* compared to the wt strain at 26°C and 37°C. The extent of differential expression was measured in terms of log2Fold Change and hyphen (-) indicates failure to meet the significance cut off or undetected. Values in red and blue indicate the log2Fold Change increases or decreases in the mutated yersinia respectively.

**Table S2**. DEGs exclusively regulated in the *m-ops* mutant compared to the wt strain at 26°C and 37°C. The extent of differential expression is measured in terms of log2Fold Change and hyphen (-) indicates failure to meet the significance cut off or undetected. Values in red and blue indicate the log2Fold Change increases or decreases in the mutated yersinia respectively.

**Table S3**. DEGs exclusively regulated in the *ΔO-aB* mutant compared to the wt strain at 26°C and 37°C. The extent of differential expression is measured in terms of log2Fold Change and hyphen (-) indicates failure to meet the significance cut off or undetected. Values in red and blue indicate the log2Fold Change increases or decreases in the mutated yersinia respectively.

**Table S4**. O-antigen operon encoding DEGs regulated in (*ΔrfaH, m-ops*, and *ΔO-aB*) mutants compared to the wt at 26°C and 37°C. The extent of differential expression is measured in terms of log2Fold Change and hyphen (-) indicates failure to meet the significance cut off or undetected. Values in red and blue indicate the log2 Fold Change increases or decreases in the mutated yersinia respectively.

**Table S5**. Strains used in this study.

**Table S6**. The primers of five differentially expressed genes used for qRT-PCR verification

## Acknowledgements

This project has received funding from the Swedish Research Council (No. 2021-02466), the Swedish Research Council Excellence Center grant (No. 2022-06543) for the Center for Modeling Adaptive Mechanisms in Living Systems Under Stress, Kempestiftelserna (JCK22-0017), and the Medical Faculty at Umeå University (FS 2.1.6-281-22) for K. Avican and UCMR Excellence by Choice’ Postdoctoral Programme in Life Science to J. K. Waititu.

## Author contributions

JKW, KN, GLM, TRDC, and KA conceived, designed, and performed the experiments. JKW, KN, GLM, TRDC, and KA analyzed the data. JKW and KA wrote the manuscript.

