## Supplementary Figure 1 for "RfaH is Essential for Virulence and Adaptive Responses in *Yersinia pseudotuberculosis* Infection"

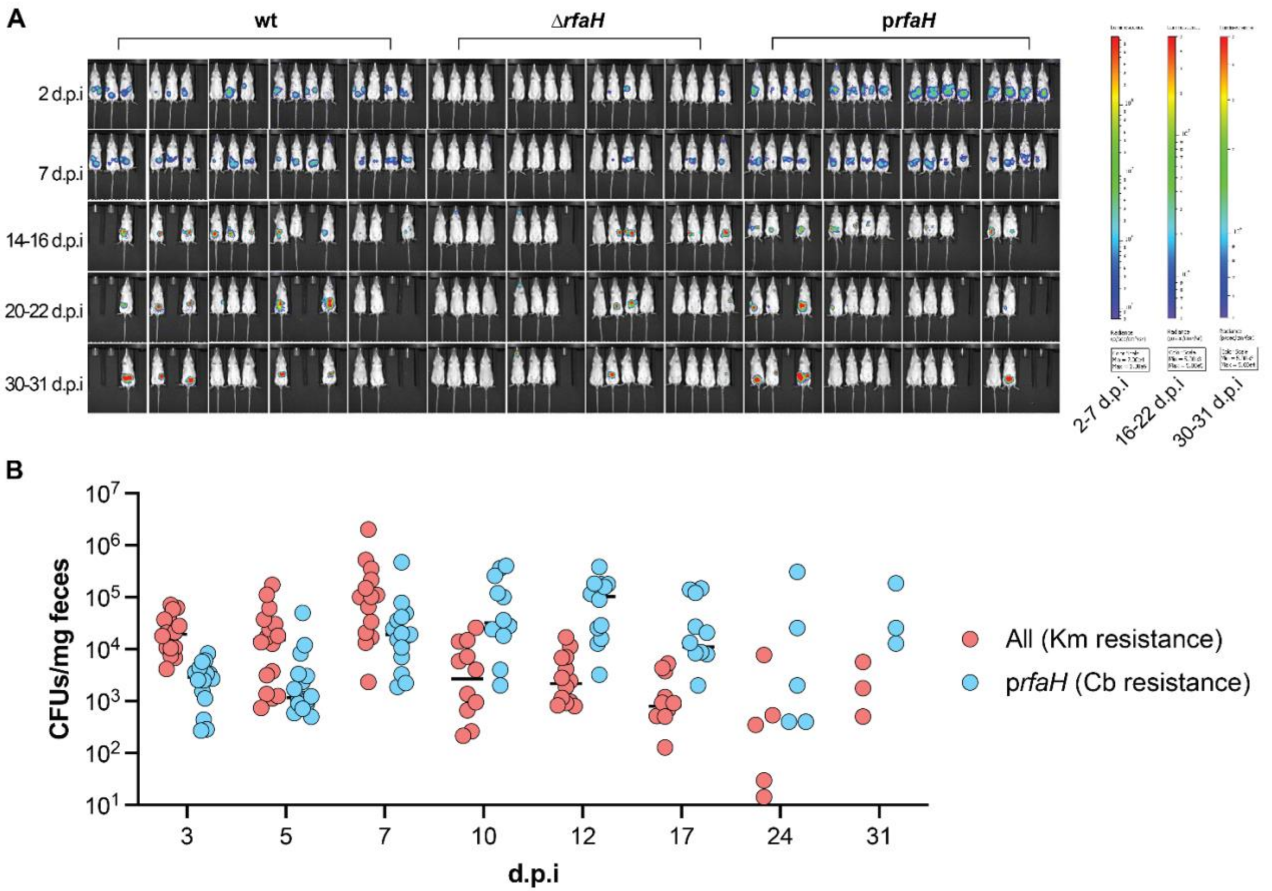


**Figure S1.** The low dose oral infection of FVB/n mouse leads to virulence attenuation in *ΔrfaH* and can be complemented with in trans expression of *rfaH*. (**A**) Mice were infected orally with overnight cultures of bacteria and the process of infection was monitored by days post-infection (d.p.i) by detecting total photon emission, using the IVIS Spectrum system. The intensity of bioluminescent emission is represented as pseudocolors with variations in color representing light intensity; red represents the most intense light emission, while blue corresponds to the weakest signal. (**B**) The screening of antibiotic resistance through monitoring the presence of *prfaH* during infection. The presence of *prfaH* was constantly monitored by screening for antibiotic resistance encoded on plasmids encoding rfaH as *Y. pseudotuberculosis* was shed through feces.
