## Supplementary Table 5 for "RfaH is Essential for Virulence and Adaptive Responses in *Yersinia pseudotuberculosis* Infection"

**Table S5**. Strains used in this study

| **Strain** | **Genotype** | **Reference** |
| --- | --- | --- |
| *E. coli* | | |
| DH5αλpir | F^−^φ80 Δ*lacZ* ΔM15 *endA1* *recA1* *hsdR17*(r_K_ ^−^ m_K_ ^+^)*supE44* *thi-1* λ^−^ *gyrA96* *relA1Δ* |  |
| S17-1λpir | *RP4-2(Km::Tn7,Tc::Mu-1), pro-*  *82, LAMpir, recA1, endA1, thiE1, hsdR17, creC510* |  |
| *Y. pseudotuberculosis* YPIII | | |
| YPIII/pIBX (originally  called Xen4) | Putative transposase (*pYV0017*)::Tn5*luxC*  *DAB*, Km^R^ | Caliper Life  Sciences, Inc. |
| YPIII, Δ*rfaH*/pIBX | Δ*rfaH* (*YPK_3937*), Km^R^ | This study |
| YPIII, *m-ops* | *TAtGgCAAGGaCaGTAtCGTGG*  *GtGGGATA*, Km^R^ | This study |
| YPIII, *ΔddhB*/pIBX | Δ*ddhB*, (*YPK_3192*), Km^R^ | This study |
| YPIII, *ΔO-aB*/pIBX | Δ*YPK_3177-3183*, Km^R^ | This study |
| YPIII, *prfaH*/pIBX | *prfaH*::*pMMB66HE*, Km^R^, Amp^R^ | This study |
